# Differential response of *Senna occidentalis* L. to arsenic and cadmium contaminated soil

**DOI:** 10.1101/2023.10.26.564171

**Authors:** Zoe Lai, Sagar Datir, Jacqueline Weber, Ebenezer Belford, Sharon Regan

**Affiliations:** Biology Department, Queen’s University, Kingston, Ontario, Canada; Naoroji Godrej Centre for Plant Research, Shirwal, India; Department of Theoretical and Applied Biology, Kwame Nkrumah University of Science and Technology, Kumasi, Ghana

**Keywords:** Antioxidant, biomass, chlorophyll, heavy metal, phytoremediation, proline

## Abstract

We investigated the phytoremediation potential of *Senna occidentalis* L., a pantropical plant that has been associated with tolerance to heavy metal-contaminated soils around mining sites. Seedlings of *S. occidentalis* were exposed to cadmium chloride (CdCl_2_) and sodium arsenate (Na_3_AsO_4_) at concentrations of 200, 300, and 400 mg L^-1^ under greenhouse conditions. Heavy metal tolerance was assessed by comparing biomass and stress indicators such as chlorophyll, proline, and hydrogen peroxide content. Arsenic treatment had more toxic effects than cadmium on *Senna* physiology. Regardless of concentration of arsenic applied, the biomass decreased by 50% as compared to control and cadmium-treated plants. Chlorophyll content decreased with exposure to both heavy metals. Higher concentration of Cd and As (400 mg L^-1^) resulted in 50% reduction in chlorophyll content. Proline and hydrogen peroxide levels were higher in arsenic-treated plants compared to controls and cadmium-treated plants, indicating an enhanced stress response when exposed to arsenic. When heavy metal content was measured, there was a significant accumulation of arsenic in the leaves, stems, and roots, indicating that arsenic in these tissues was responsible for the profound changes in biomass, proline, and hydrogen peroxide content. In contrast, although significant, there was less cadmium uptake by *Senna* and tolerance can be seen, which was reflected by normal biomass, proline, and hydrogen peroxide levels. High translocation of metals from soil into roots and low translocation from root to shoot tissues suggests the potential for *S. occidentalis* to be used for phytostabilization of arsenic- and cadmium-contaminated soils.

## Introduction

Worldwide, there are about 5 million sites contaminated with a wide variety of toxic heavy metals (∼20 million ha of land) (Wuana and Okieimen 2011; He et al. 2015). The impact on people is especially high in areas surrounding mines, such as in the Giant Mine in Yellowknife, NWT, Canada (Jamieson 2014), and around gold mines in Obusai, Ghana (Bempah and Ewusi 2016). In Ghana and elsewhere, farming occurs near these mines (Essumang et al. 2007), and arsenic (As), cadmium (Cd), and other contaminants have been observed in these soils which can then be absorbed by crops (Essumang et al. 2007; Bempah and Ewusi 2016). Both Cd and As are highly harmful contaminants in the environment that lead to heavy metal pollution of soil and water due to natural weathering of geological materials, industrialization, pesticides, fertilizers, mining, and manufacturing (Garelick et al. 2009; Govindasamy et al. 2011; Ozbek et al. 2014; Amel et al. 2016). Due to their ability to accumulate through the food chain, Cd and As can adversely affect humans (Essumang et al. 2007; Reeves and Chaney 2008; Huang et al. 2017; Shahid et al. 2017) where chronic Cd and As exposure has been connected to cancer and several other diseases (Reeves and Chaney 2008; Järup and Akesson 2009; Jomova et al. 2011; Kamunda et al. 2016; Huang et al. 2017; Shahid et al. 2017; Ayangbenro and Babalola 2017). Cd and As exposure occur through the consumption of contaminated crops grown on polluted soil and As is also consumed by drinking contaminated water. Since these heavy metals can persist indefinitely in the environment, and cannot be chemically removed nor biodegraded, they must be physically removed or transformed into non-toxic compounds (Garbisu and Alkorta 2001).

*Senna occidentalis* L (formerly *Cassia occidentalis*) is an abundantly growing pantropical medicinally important plant that has been identified as a hyperaccumulator of heavy metals (Yadav et al 2010; Juliana et al. 2019). It has a large biomass, has shown tolerance to high levels of heavy metal exposure, and grows well in areas known to have Cd . Although *Senna occidentalis* has been previously studied as a potential phytoremediator plant with great promise (Zauro et al. 2015; Kulshrestha and Dabra 2016; Juliana et al. 2019), no previous study has characterized the individual effects of Cd and As on the tolerance ability and physiological characteristics. Silva et al. (2018) studied the effects of Cd on *Cassia alata* plants and based on increased antioxidant enzyme activities in Cd-treated plants compared to control plants, they concluded that *C. alata* could be used in phytoremediation programs. They suggested that efficient activity of antioxidant enzymes helped minimize the oxidative stress caused by Cd exposure and consequently improved the protection of metabolic pathways. Biochemical indicators might help in determining the degree of contamination of the contaminated sites exposed to heavy metals (Petrovic and Krivokapic 2020). Juliana et al. (2019) have investigated the phytoextraction capability of *Senna occidentalis* by examining its uptake of Cd, Cu, Ni, Pb and Zn from waste dump soil site in Nigeria. They concluded that *S. occidentalis* was capable of taking up all five heavy metals by accumulating and translocating to the aboveground harvestable biomass.

In this study we investigated the effect of varying levels of Cd and As on *S. occidentalis* physiology under greenhouse conditions. The aim of this study is to identify the biochemical and physiological parameters that could explain the uptake, survival strategy and sensitivity to metal toxicity of *S. occidentalis* for individual heavy metal. The phytoremediation potential of *S. occidentalis* was determined based on the metal toxicity indicators such as photosynthetic pigments, proline, and antioxidant enzyme levels. This study also determined the extent of Cd and As accumulation in different tissues, and to will help identify the factors that contribute to the tolerance mechanism of *S. occidentalis* under Cd and As toxicity.

## Materials and Methods

### Plant material and growth conditions

Seeds of *Senna occidentalis* were collected from wasteland in Obuasi, Ghana, a mining community that is rich in gold deposits. A greenhouse experiment was carried out in the Phytotron facility at Queen’s University Biosciences Complex. Dry seeds were first rooted in a 50-50 sand-soil mixture (ASB Greenworld, Original Grower Mix, GM 15% with perlite) for one month before transplanting into ASB Greenworld soil in individual pots. A total of 32 6-inch-wide x 4.5-inch-tall pots with saucers, containing one plant per pot, were grown in a randomized arrangement in four replications at approximately 25°C with natural and supplemental lighting (16 hr daylight). Cadmium chloride (CdCl_2_), sodium arsenate (Na_3_AsO_4_), and water (pH 7.0; control) were individually added to pots at concentrations of 200, 300, and 400 mg L^-1^ twice a week (100 mL each solution ensuring that the solution does not leak from the pot) for 4 weeks (total 8 times). Leaf, stem and/or root tissues were collected from each plant after 75 days of vegetative growth and stored at –80°C for biochemical analysis.

### Biomass assessment

Leaf, stem, and root tissues were individually collected from four different plants from each treatment, and soil samples were collected from each pot. Roots were thoroughly washed to remove all soil, and leaves, stems and roots from each plant were dried in a hot air oven (65°C for two weeks) before the dry biomass was determined.

### Chlorophyll content

Chlorophyll (chlorophyll-a, chlorophyll-b, and total chlorophyll) content was determined according to Arnon (1949). Leaf tissue (0.1 g) was ground in liquid nitrogen and the chlorophyll were extracted in 2 mL of 80 % acetone (Fisher Chemical, ON, Canada). Samples were centrifuged at 2739 x g at 4 °C for 15 min. The absorbance of the supernatant was measured at 663 and 645 nm using acetone as a blank. The chlorophyll content was expressed in mg g^-1^ FW^-1^ (fresh weight). The amount of chlorophyll present in the extract was calculated using the following formula:

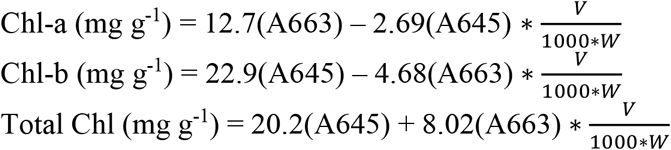

where, A = absorbance at specific wavelengths, V = volume of 80% acetone and W = fresh weight of tissue.

### Proline content

Proline content was determined as described by Bates et al. (1973). Flash-frozen plant samples (0.2 g) were ground and extracted using 2 mL of 3 % 5-sulfosalicylic acid (Sigma, ON, Canada). The extract was centrifuged for 10 min at 8050 x g at 4 °C. A reaction mixture of 100 μL of supernatant, 900 μL of 3 % sulfosalicylic acid, 2 mL of glacial acetic acid, and 2 mL of acid ninhydrin (glacial acetic acid, 6 M phosphoric acid, ninhydrin (2,2-dihydroxyindane-1,3-dione, Sigma, ON, Canada) were boiled in a water bath at 100 °C for 1 hr. Proline was extracted with 4 mL toluene (Fisher Chemical, ON, Canada) and the absorbance of the toluene layer was measured at 520 nm. Reference standards of proline (0 to 40 μg mL^-1^) (Sigma, ON, Canada) were prepared in 3 % 5-sulfosalicylic acid and the amount of proline was expressed as μmoles proline g^-1^ FW^-1^ (fresh weight).

### Hydrogen peroxide content

Hydrogen peroxide (H_2_O_2_) content was determined according to Velikova et al. (2000). Flash-frozen plant tissue (0.2 g) were ground and extracted in 2 mL of trichloroacetic acid (0.1 % w/v). The extract was centrifuged for 15 min at 8050 x g at 4 °C. A mixture of 0.5 mL of supernatant, 0.5 mL of 10 mM potassium phosphate buffer (pH 7.0), and 1 mL of 1M potassium iodide as vortexed briefly and the absorbance was measured at 390 nm. A mixture of potassium phosphate buffer, potassium iodide, and trichloroacetic acid was used as the blank. H_2_O_2_ content was quantified by referencing a calibration curve made with a 0 to 200 μM concentration range and expressed as μmole H_2_O_2_ g^-1^ FW^-1^ (fresh weight).

### Arsenic and cadmium content in plant tissues

Leaf, stem, and root samples were homogenized separately in a coffee grinder (Proctor Silex Fresh Grind™) and sent to the Queen’s Biosciences Analytical Services Unit for Inductively Coupled Plasma Mass Spectrometry (ICP-MS) to determine the As and Cd contents of the individual plant tissues (leaf, shoot, and root). From each sample, 0.5 g of ground material was reduced to ash in a blast furnace and digested in an acid bath consisting of nitric acid and hydrochloric acid (method revised from U.S. EPA, 2001). Analysis of the elemental composition of these samples was performed using 7700x ICP-MS, Agilent Technologies. The analytical standard for ICP-MS analyses was Spinach 1570a Certified Reference Material. Scandium, indium, and bismuth were used as internal standards.

### Statistical analysis

The plants were arranged in a random block design. All the experimental analyses except for heavy metal concentrations were performed with four replicate samples while heavy metal concentration analysis was performed on three replicates. The statistical analyses were performed on Microsoft Excel for Mac (Version 16.16.5) and RStudio Version 1.2.5033 (R version 3.6.3). Datasets underwent Levene’s Test to evaluate the homogeneity of variance as well as Shapiro-Wilks Test to examine if the data was normally distributed. Since the data did not violate these assumptions, one-way ANOVA was then performed on all datasets. A significance level of P < 0.05 and a confidence level of 95 % was used for all analyses. Error bars in graphs represent ± standard error (SE).

## Results

### Arsenic decreases plant biomass of *Senna occidentalis*

To determine the impact of heavy metal exposure on plant health, the dry biomass of the leaves, stem, and roots were measured from the heavy metal-stressed plants and compared to the control-grown plants (Fig. 1A). When comparing As-treated and Cd-treated plants, there were significantly less leaf, stem, root, and total dry biomass (leaf + stem + root) with As treatment compared to Cd-treated and control plants. Plants exposed to 200, 300 or 400 mg L^-1^ As exhibited a 41-56% decrease in leaf biomass (Fig. 1Ai) while stem biomass decreased by 36-54% as compared to the control plants (Fig. 1Aii). However, the largest biomass reduction was noticed in root samples with a 53-66% decrease in biomass (Fig. 1Aiii).

**Figure 1.**
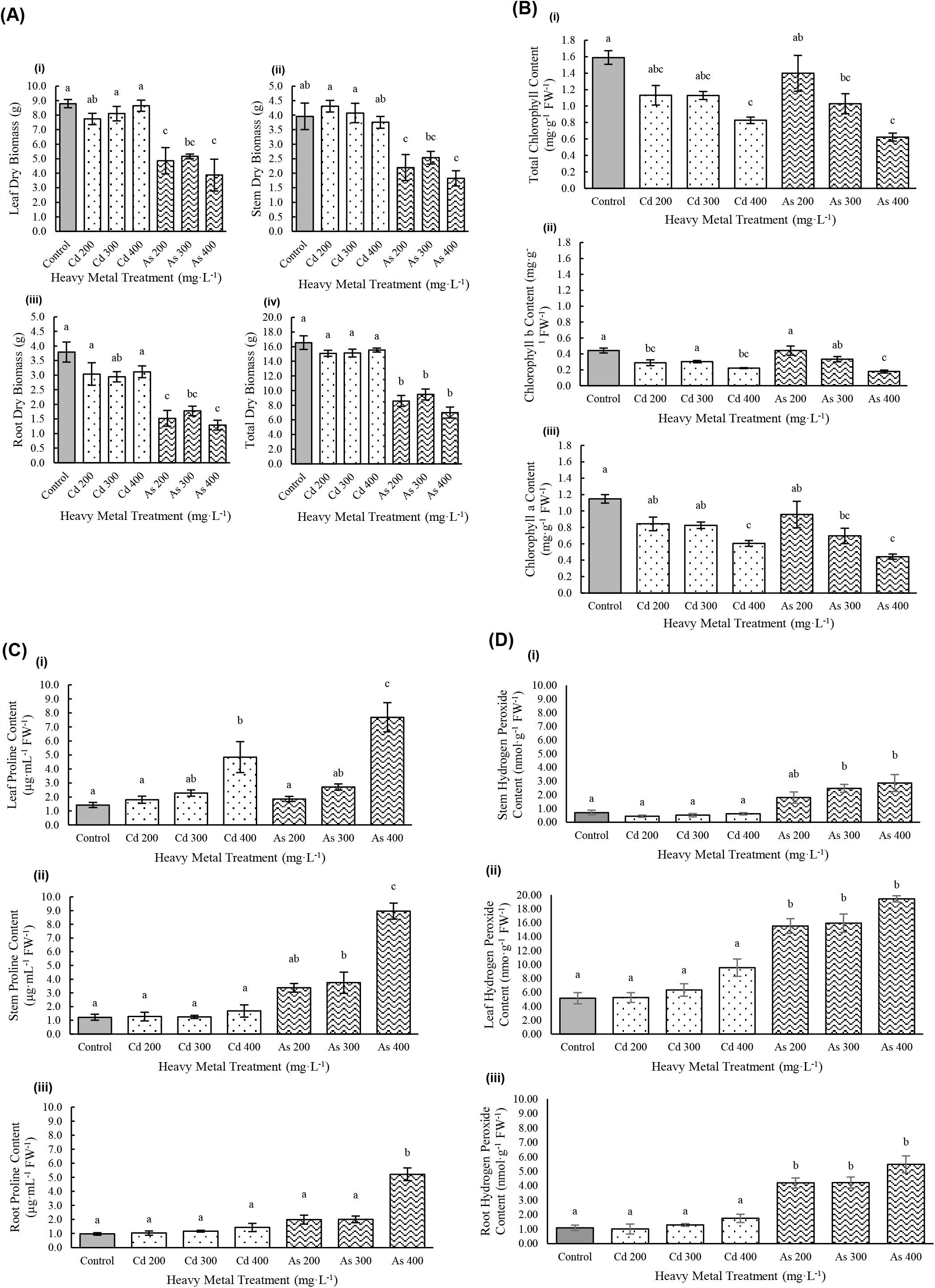
Effect of heavy metal treatments (Cd or As) on biological and physiological parameters. (A) Effect of heavy metal treatments (Cd or As) on (i) leaf, (ii) stem, (iii) root, and (iv) total biomass (g) of *S. occidentalis* (*n* = 4). (B) Effect of Cd and As heavy metal treatments on (i) total chlorophyll (ii) chlorophyll b, and (iii) chlorophyll a content (mg g^-1^ FW^-1^) in *S. occidentalis* (*n* = 4). (C) Effect of Cd and As heavy metal treatments on proline content (μg mL^-1^ FW^-1^) in the (i) leaves, (ii) stem, and (iii) roots of *S. occidentalis* (*n =* 4). (D) Effect of Cd and As heavy metal treatments on hydrogen peroxide content (nmol g^-1^ FW^-1^) in the (i) stem, (ii) leaf, and (iii) roots of *S. occidentalis* (*n =* 4). Vertical bars indicate mean ± standard error. Different letters represent significant difference between the treatments at p < 0.05.

### Chlorophyll content decreases with increasing heavy metal exposure

Chlorophyll content was analysed in all plant tissues as a potential measure of plant health. In all chlorophyll measurements (chlorophyll-a, chlorophyll-b, and total chlorophyll), there was a decrease in chlorophyll content as both As and Cd heavy metal treatments increased (Fig. 1B). The chlorophyll-a content experienced a 26-47% decrease as Cd treatments increased from 200 to 400 mg L^-1^, and a 17-61% decrease as As treatments increased (Fig. 1Biii). The most significant differences were from the highest heavy metal treatment, Cd 400 mg L^-1^ and As 400 mg L^-1^, a 47% and 61% decrease respectively compared to control. Chlorophyll-b content experienced a similar trend, and as Cd treatments increased in concentration, chlorophyll-b content decreased significantly by 35% and 50% at 200 and 400 mg L^-1^, respectively (Fig. 1Bii). As the treatments for As increased to 400 mg L^-1^, there was a significant 60% decrease in chlorophyll-b content. Thus, total chlorophyll content reflected these trends and plants treated with 400 mg L^-1^ of Cd or As experienced a significant decrease of 48% and 61% of total chlorophyll respectively compared to the control treatment (Fig. 1Bi).

### Proline accumulates in arsenic-treated plants

Proline content was measured to assess the extent of heavy metal stress of *Senna* plants in leaf, stem, and root (Fig. 1C). There was a consistent trend of increasing proline content as heavy metal treatments increased across all tissues. Significantly increased (∼3-fold) proline content was observed in leaf tissues of those treated with 400 mg L^-1^ of Cd in comparison to control (Fig. 1Ci). In the As-treated plants, as the treatment concentrations increased, so did proline content, with the highest treatment of As 400 mg L^-1^ containing significantly higher proline content by ∼5-fold compared to the control amounts. The stem tissues of As-treated plants experienced a significant increase in proline compared to control and Cd-treated plants, specifically with 300 and 400 mg L^-1^ of As that increased by ∼3-fold and ∼7-fold respectively (Fig. 1Bii). Among the root samples, only the tissues from plants treated with As 400 mg L^-1^ accumulated significantly more proline by ∼5-fold more than control (Fig. 1Biii).

### Hydrogen peroxide accumulates in arsenic-treated plants

H_2_O_2_ was used as another measure of the impact of heavy metal exposure on plant health (Fig. 1D). A trend of increasing H_2_O_2_ can be seen as heavy metal treatments increased in all plant tissues, however there were significantly increased amounts only in As treatments. The H_2_O_2_ content was significantly highest in leaf tissues and experienced more than a 3-fold increase across all As-treatments in comparison to the control leaf content (Fig. 1Dii). The stem tissues of those treated with As 300 and As 400 experienced about a significant ∼4-fold increase in H_2_O_2_ content in comparison to the control stem tissues (Fig. 1Di). Root tissues experienced the significantly ∼5-fold increase H_2_O_2_ content between control and As-treated plants (Fig. 1Diii).

### Cadmium and arsenic accumulation in plant tissues

The concentration of Cd and As was measured in the Cd- and As-treated as well as in control plants of *S. occidentalis* (Fig. 2). For both the Cd- and As-treatment groups, plants accumulated the highest concentration of heavy metals in the roots, followed by stem, and then leaves. In Cd-treated plants, Cd levels were significantly higher in roots, stems, and leaves than control plants, but these levels were approximately 5 to 14-fold lower than the amount of As that accumulated in the tissues of the As-treated plants. The highest concentration of As was found in the roots at an average of 570 μg g^-1^ while the stems and leaves accumulated approximately 50 μg g^-1^ and 25 μg g^-1^ respectively amongst all As treatments (200, 300, and 400 mg L^-1^) (Fig. 2C). Meanwhile, the root, stem, and leaf tissues of the Cd-treated plants averaged approximately 100 μg g^-1^, 7 μg g^-1^, and 2 μg g^-1^ of Cd respectively across all the treatments of Cd (200, 300, and 400 mg L^-1^) (Fig. 2).

**Figure 2.**
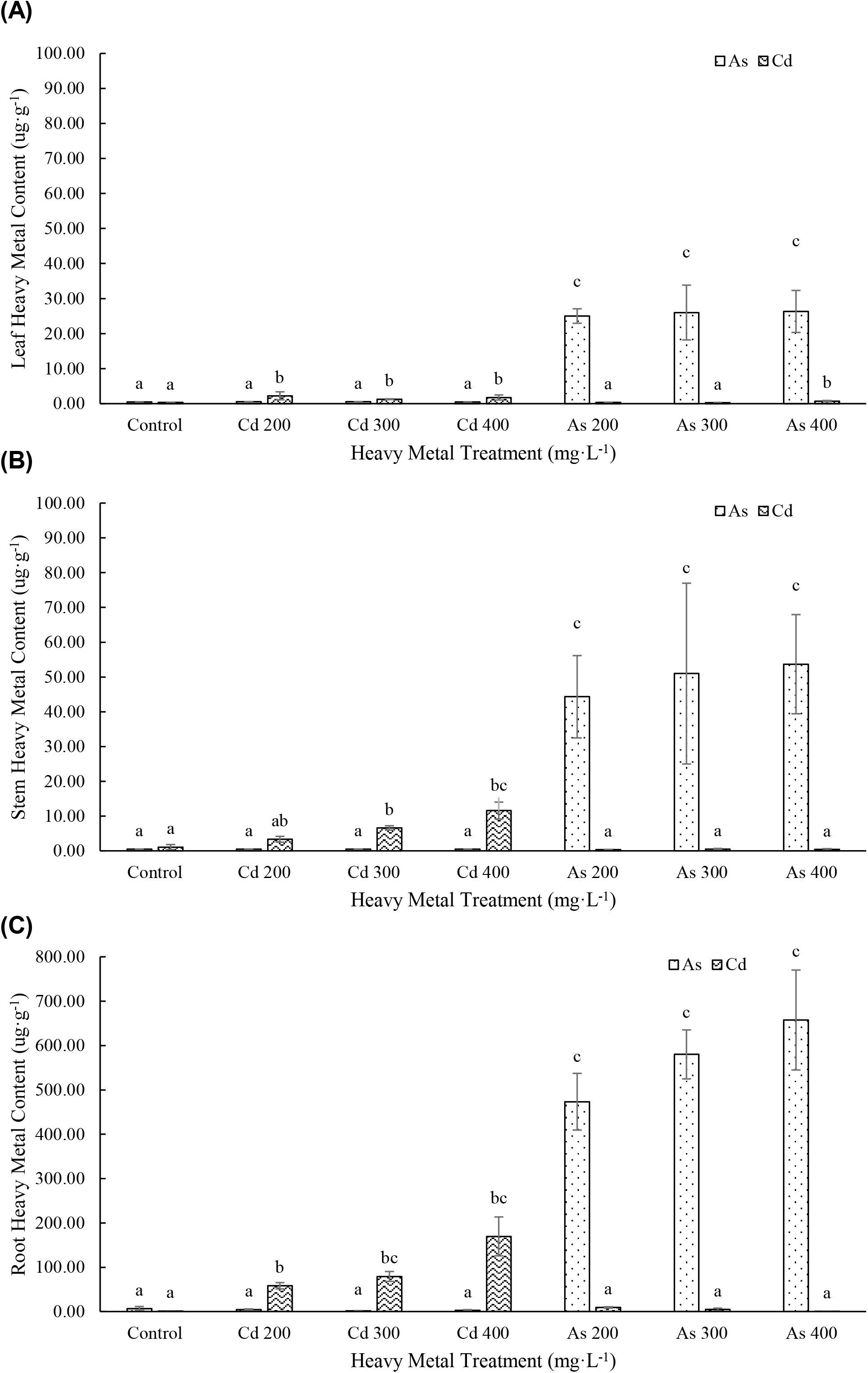
Heavy metal concentration in the (A) leaf, (B) stem, and (C) root tissues of *S. occidentalis* exposed to a range of Cd and As treatments (*n =* 3). Vertical bars indicate mean ± standard error. Different letters represent significant difference between the treatments at P < 0.05. Note the 8 times y-axis scaling between (C) and (A), (B).

### Assessment of phytoremediation potential of Senna occidentalis

Phytoremediation potential was evaluated based on the translocation factor (TF), bioconcentration factor (BCF), bioaccumulation coefficient (BAC), and concentration index (CI) (Marchiol *et al*. 2004; Ghosh and Singh 2005; Yoon *et al*. 2006; Chandra and Hoduck 2018). TF can be used to measure the plant’s ability to translocate metals from the roots to the shoots and was determined by a ratio of metal concentration in the shoots and roots. BCF can determine the ability of the plant to uptake heavy metals from the soil into the roots and was calculated as the ratio of metals in the roots and the soil. BAC measured heavy metal movement from the soil into the shoot of the plant and was calculated as the ratio of the metal concentration in the shoot and soil. CI was calculated as the ratio of heavy metal concentration in experimental plants compared to the control and can assess the metal accumulation in the plant. Combined, these factors can be used to assess the potential of a plant species for phytoremediation purposes.

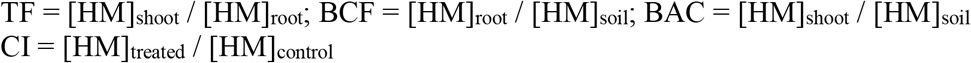

The phytoremediation potential was determined based on the tendency of the plants to accumulate the heavy metals in tissues. In the case of TF values, both Cd and As treatments resulted in TF values less than 1.0 (Table 1). The BCF values were greater than 1.0 for all Cd and As heavy metal treatments (Table 1). On the other hand, the BAC values were less than 1.0 across all treatments, except for the plants treated with 200 mg L^-1^ of As (Table 1). For both heavy metal treatments, the CI values increased with increasing heavy metal treatment concentrations. The CI values were higher in the As-treated plants as compared to Cd-treated, with the highest values at the 400 mg L^-1^ heavy metal treatment, 96.66 and 161.66 for Cd and As respectively (Table 1).

**Table 1.** The translocation factor (TF), bioconcentration factor (BCF), biological accumulation coefficient (BAC), and concentration index (CI) of *Senna occidentalis* under three concentrations of Cd and As treatments. The values are mean ± standard error (*n* = 3).

| Heavy metal<br>concentration<br>(mg·L<sup>-1</sup>) | TF |  | BCF |  | BAC |  | CI |  |
| --- | --- | --- | --- | --- | --- | --- | --- | --- |
|  | Cd | As | Cd | As | Cd | As | Cd | As |
| 200 | 0.15 ± | 0.16 ± | 1.10 ± | 9.34 ± | 0.17 ± | 1.28 ± | 28.24 ± | 125.69 ± |
|  | 0.03 | 0.05 | 0.15 | 2.00 | 0.04 | 0.13 | 7.78 | 54.7 |
| 300 | 0.10 ± | 0.14 ± | 3.42 ± | 3.51 ± | 0.39 ± | 0.52 ± | 42.89 ± | 156.13 ± |
|  | 0.01 | 0.04 | 2.06 | 0.51 | 0.27 | 0.23 | 14.84 | 66.04 |
| 400 | 0.09 ± | 0.14 ± | 7.93 ± | 6.87 ± | 0.68 ± | 0.94 ± | 96.66 ± | 161.66 ± |
|  | 0.04 | 0.04 | 2.90 | 1.41 | 0.29 | 0.27 | 45.03 | 58.61 |

## Discussion

Heavy metal contamination of soil and water are of increasing concern worldwide and require more sustainable methods for cleanup, and phytoremediation offers a cost-effective mechanism. In this study, we assessed the ability of *S. occidentalis* to uptake Cd and As from the soil. The impact of these metals on plant growth and physiology of *Senna* was assessed through analysis of dry biomass as well as quantification of chlorophyll, proline, and H_2_O_2_ content in tissues. Previous studies have suggested that the growth performance of the heavy metal-treated plants can be assessed based on the impact on biomass, and this can be utilized for the determination of the phytoremediation potentiality of plants (Ullah et al. 2020). In this study we found that As-treated plants had significantly decreased leaf, stem, root and total biomass, in comparison to Cd-treated and control plants (Fig. 1A). Cd-treated plants appeared unaffected suggesting poor mobilization of Cd in *Senna occidentalis* plants (Godinho et al. 2018). The decreased biomass in As-treated plants could be attributed to disturbances in cellular metabolic processes and essential nutrients absorption and distribution (Shanker et al. 2005; Ullah et al. 2020). While As treatments exhibited almost 50 % reduction in the total dry mass, plant survival under various As concentrations, especially under 400 mg L^-1^, suggest its tolerance of As (Fig. 1A). In agreement with previous studies, As toxicity induced low biomass in crops attributed to high metal concentrations in the shoots and roots (Liu et al. 2007; Niazi et al. 2017; Ullah et al. 2020). Biomass is an important parameter to examine when assessing plants stress responses, as it provides insight into the negative impact on the plant’s health (Mukhtar et al. 2010; Tiwari et al. 2016). Additionally, it is an important trait to consider when examining a plant species for phytoremediation since high biomass would allow for more accumulation of heavy metals than a plant with equal translocation of metals into its tissues but lower biomass.

Chlorophyll quantification can be used to assess general plant health of heavy metal-treated plants. It measures the plant’s photosynthetic capabilities and any damages to the photosynthetic system due to heavy metal stress can be seen from chlorophyll quantification (Pietrini et al. 2003; Mrnka et al. 2012). Cd and As exposure in plants have been found to disrupt chloroplast membranes and damage photosynthetic enzymes or pigments, leading to impairment of photosynthetic capabilities, ultimately resulting in decreased chlorophyll content (Sánchez-Viveros et al. 2011; Marmiroli et al. 2013; Tiwari et al. 2016; Zhang et al. 2020). *S. occidentalis* was found to show decreased chlorophyll content as heavy metal concentrations increased, with significant decrease of ∼50 % all chlorophyll content at 400 mg L^-1^ Cd and As concentration (Fig. 1B).

The increase in proline content as heavy metal concentrations increased supports the role of proline in maintaining plant growth during heavy metal stress, as it is an important osmolyte (Verbruggen and Hermans 2008; Irfan et al. 2014; Hashem et al. 2016). Proline is involved with membrane protection, the neutralization of toxic reactive oxygen species, and it may protect some specific enzymes from being damaged (Handique and Handique 2009; Irfan et al. 2014; Hashem et al. 2016). The only significant difference between the proline content of the control plants to the tissues of the Cd-treated plants where in the leaves of those treated with 400 mg L^-1^ (Fig. 1C), indicating that *Senna* plants did not exhibit signs of Cd stress until exposed to high levels. There was, however, significantly higher proline in shoots (leaf + stem) compared to roots in As-treated plants. Accumulation of more proline in shoots has been attributed to maintenance of chlorophyll level and cell turgor to protect the photosynthetic activity under salt stress (Silva-Ortega et al. 2008). In nickel-treated pea plants, it was found that having lower root proline content compared to leaves could protect leaf tissues against dehydration, since water is a crucial component for photosynthesis reactions (Gajewska and Skłodowska 2005). The accumulation of proline may help in the detoxification of As, as well as reduce uptake and damage to membranes and proteins (Wu et al. 1998; Siripornadulsil et al. 2002; Mishra and Dubey 2006). The increase in proline levels in leaf, stem and root tissues indicates the physiological response of *Senna* plants under As stress, suggesting an antioxidant role in detoxification of heavy metal accumulation as seen in other plant species (Singh and Sinha 2005; Atabaki et al. 2020).

Plants are known to accumulate higher quantities of hydrogen peroxide (H_2_O_2_) as both a defensive measure against heavy metal damage and a direct result of the damage in the form of oxidative stress (Singh et al. 2007; Opdenakker et al. 2012; Huang et al. 2020). The trend of increasing H_2_O_2_ content in *S. occidentalis* followed by increasing heavy metal treatment demonstrates the enhanced production of H_2_O_2_ during heavy metal stress. Our study found that there was a significantly higher accumulation of H_2_O_2_ by more than 3-fold in As treatments than in control plants (Fig. 1D). The differential increase in H_2_O_2_ content in As treatments indicate the possibility that H_2_O_2_ acted as a defense molecule in heavy metal tolerance (Singh et al. 2007; Asgher et al. 2021). The significant increase in H_2_O_2_ content in all plant tissues with As exposure (Fig. 1D) may be due to oxidative stress caused by As toxicity. Exposure to As has been found to generate ROS, increased H_2_O_2_ content in plant tissues, and result in lipid peroxidation, cellular damage, and cell death (Hartley-Whitaker et al. 2001; Ahsan et al. 2008; Farooq et al. 2015; Majumder et al. 2018).

The analysis of heavy metal content in different tissues revealed that the highest Cd and As concentrations were observed in root followed by stem and leaves, indicating poor mobilization of metals into aerial tissues from the roots (Fig. 2). The low concentration of Cd in plant tissues indicates that *S. occidentalis* was able to exclude most Cd from entering the aerial parts of the plant. Other plants have developed protection mechanism from Cd through avoidance or exclusion. *Nicotiana tabacum* have been shown to excrete cadmium-calcium crystals in the heads of its trichomes (Choi et al. 2001). Martin et al. (2012) also found natural Cd-specific exclusion in a *Thlaspi arvense* ecotype that was efficient in excluding Cd from its roots and shoots tissues. At this time, we do not know how *S. occidentalis* is limiting the uptake and/or accumulation of Cd in its tissues. In contrast to the levels of Cd, *S. occidentalis* accumulated up to 5-fold more As in the roots, stems and leaves (Fig. 2). These high levels of As throughout the plant are likely responsible for the low biomass (Fig. 1A), high proline levels (Fig. 1C), and high H_2_O_2_ levels (Fig. 1D) in these plants. While arsenate is generally readily transported into plant roots through phosphate transporters because the chemical structure is analogous to Pi (Meharg and Macnaib 1990), at this time, we do not know the specific mechanism behind As uptake in *S. occidentalis*. Irrespective of the impact of As on plant growth, *S. occidentalis* can uptake significant amounts of As and could be a viable candidate for phytoremediation.

Phytoremediation potential of plants can be assessed based on the TF, BCF, BAC, and CI values (Table 1). These factors help to identify the suitability of plants for phytoremediation (i.e., phytoextraction or phytostabilization) by explaining the accumulation characteristics and translocation behaviours of metals in plants (Mellem et al. 2012; Amin et al. 2018). The TF is an indication of the ability of the plant to translocate metals from the roots to the aerial parts of the plant (Marchiol et al. 2004), while the BCF and BAC is an index of the ability of the plant to accumulate a particular metal with respect to its concentration in the soil into the roots or aboveground tissues respectively (Ghosh and Singh 2005). CI provides more detail on the plant’s ability to accumulate the metals into the whole plant compared to control plants. *S. occidentalis* has an average TF value < 1, BCF value > 1, BAC value < 1, and high CI values for both Cd and As treatments (Table 1). Plants with a high BCF value and a low TF value has the potential to be used for phytostabilization of contaminated soils (Yoon et al. 2006). Therefore, it can be concluded that *S. occidentalis* is a good phytostabilizer for As, with potential in Cd phytostabilization, due to its heavy metal tolerance and accumulation pattern. The Cd accumulation was shown to mainly be restricted to the root tissue, supplemented by a lower translocation rate to shoots (Dai et al. 2011, Fan et al. 2011; Silva et al. 2018). These results are in conformation with Kulshrestha and Dabral (2016) who tested absorption and their translocation of Cd from the soil in *Cassia occidentalis*. They concluded that *Cassia occidentalis* showed moderate absorption and translocation of Cd from roots to above ground parts. Overall, *S. occidentalis* shows low levels of Cd absorption and has the ability to accumulate high levels of As.

## Conclusions

Here we show that *S. occidentalis* can accumulate more As in comparison to Cd in their roots, stems, and leaves. Using a selection of physiological assays, we find that dry biomass, proline, and H_2_O_2_ content revealed that the plants are under significant stress when treated with As but are not obviously impacted by Cd. These parameters can be used as stress indicators to explain the uptake, survival strategy and sensitivity to metal toxicity of *S. occidentalis*. This study reveals the potential of using these simple assays to identify contaminated sites in a costeffective way. Additionally, the accumulation of As in *S. occidentalis* tissues presents this plant as a suitable candidate for the stabilization of the heavy metal in contaminated soil. Field-based research is required to evaluate the significance of stress indicators in determining the metal tolerance and phytoremediation potential of *S. occidentalis*.

## Author Contributions

EB collected the seeds and helped to design the study with SR. ZL conducted the greenhouse trial and the experimental analysis under the guidance of SR and SD. JW and SD carried out preliminary analyses to develop the protocols for this study. ZL and SD analyzed the data and drafted the manuscript. EB and SR did the critical revision of the manuscript.

## Acknowledgements

The authors would like to thank McKaila Anderson for technical assistance.

## References

Ahsan N, Lee DG, Alam I, Kim PJ, Lee JJ, Ahn YO, Kwak SS, Lee IJ, Bahk JD, Kang KY, et al. (2008) Comparative proteomic study of arsenic-induced differentially expressed proteins in rice roots reveals glutathione plays a central role during As stress. Proteomics 8:3561–3576. 10.1002/pmic.200701189

Amel SB, Nabil M, Nadia A, Hocine G, Hakim L, Nadjib D (2016) Phytoremediation of soil contaminated with Zn using Canola (Brassica napus L.). Ecol Eng 95:43–49. 10.1016/j.ecoleng.2016.06.064

Amin H, Arain BA, Jahangir TM, Abbasi MS, Amin F (2018) Accumulation and distribution of lead (Pb) in plant tissues of guar (Cyamopsis tetragonoloba L.) and sesame (Sesamum indicum L.): Profitable phytoremediation with biofuel crops. Geol Ecol Landsc 2:51–60. 10.1080/24749508.2018.1452464

Arnon DI (1949) Copper enzymes in isolated chloroplasts, polyphenol oxidase in Beta vulgaris. J Plant Physiol 24:1–15.

Asgher M, Ahmed S, Sehar Z, Gautam H, Gandhi SG, Khan NA (2021) Hydrogen peroxide modulates activity and expression of antioxidant enzymes and protects photosynthetic activity from arsenic damage in rice (Oryza sativa L.). J Hazard Mater 401:123365. doi: 10.1016/j.jhazmat.2020.123365.

Atabaki N, Shaharuddin NA, Ahmad SA, Nulit R, Abiri R (2020) Assessment of water Mimosa (Neptunia oleracea Lour.) Morphological, physiological, and removal efficiency for phytoremediation of Arsenic-polluted water. Plants 9:1500. 10.3390/plants9111500

Ayangbenro AS, Babalola OO (2017) A new strategy for heavy metal polluted environments: A review of microbial biosorbents. Int J Environ Res Public Health 14:1–16. 10.3390/ijerph14010094

Bates LS (1973) Rapid determination of free proline for water stress studies. Plant Soil 39:205–207.

Bempah CK, Ewusi A (2016) Heavy metals contamination and human health risk assessment around Obuasi gold mine in Ghana. Environ Monit Assess 188:261. 10.1007/s10661-016-5241-3

Chandra R, Hoduck K (2018) Phytoremediation and Physiological Effects of Mixed Heavy Metals on Poplar Hybrids. Heavy Metals. London (UK): IntechOpen. 10.5772/intechopen.76348

Choi YE, Harada E, Wada M, Tsuboi H, Morita Y, Kusano T, Sano H (2001) Detoxification of cadmium in tobacco plants: formation and active excretion of crystals containing cadmium and calcium through trichomes. Planta 213:45–50. 10.1007/s004250000487

Dai ZY, Shu WS, Liao B, Wan CY, Li JT (2011) Intraspecific variation in cadmium tolerance and accumulation of a high biomass tropical tree Averrhoa carambola L.: implication for phytoextraction. J Environ Monit 13:1723–1729. 10.1039/c1em10054h

Essumang DK, Dodoo DK, Obiri S, Yaney JY (2007) Arsenic, cadmium, and mercury in cocoyam (Xanthosoma sagititolium) and watercocoyam (Colocasia esculenta) in Tarkwa a mining community. B Environ Contam Tox 79:377–379. 10.1007/s00128-007-9244-1

Fan KC, His HC, Chen CW, Lee HL, Hseu ZY (2011) Cadmium accumulation and tolerance of mahogany (Swietenia macrophylla) seedlings for phytoextraction applications. J Environ Manage 92:2818–2822. 10.1016/j.jenvman.2011.06.032

Farooq MA, Gill RA, Ali B, Wang J, Islam F, Ali S, Zhou W (2015) Subcellular distribution, modulation of antioxidant and stress-related genes response to arsenic in Brassica napus L. Ecotoxicology 25:350–366. 10.1007/s10646-015-1594-6

Gajewska E, Skłodowska M (2008) Differential biochemical responses of wheat shoots and roots to nickel stress: Antioxidative reactions and proline accumulation. Plant Growth Regul 54:179–188. 10.1007/s10725-007-9240-9

Garbisu C, Alkorta I (2001) Phytoextraction: a cost-effective plant-based technology for the removal of metals from the environment. Bioresour Technol 77:229–236. 10.1016/S0960-8524(00)00108-5

Garelick H, Jones H, Dybowska A, Valsami-Jones E (2009). Arsenic Pollution Sources. In: Reviews of Environmental Contamination. Vol. 197. New York (NY): Springer. p. 17–60. 10.1007/978-0-387-79284-2_2

Ghosh M, Singh SP (2005) A comparative study of cadmium phytoextraction by accumulator and weed species. Environ Poll 133:365–371. 10.1016/j.envpol.2004.05.015

Godinho DP, Serrano HC, Da Silva AB, Branquinho C, Magalhães S (2018) Effect of cadmium accumulation on the performance of plants and of herbivores that cope differently with organic defenses. Front Plant Sci 9:1723. 10.3389/fpls.2018.01723

Govindasamy C, Arulpriya M, Ruban P, Francisca LJ, Ilayaraja A (2011) Concentration of heavy metals in seagrasses tissue of the Palk Strait, Bay of Bengal. Int J Environ Sci 2:145–153.

Handique GK, Handique AK (2009) Proline accumulation in lemongrass (Cymbopogon flexuosus Stapf.) due to heavy metal stress. J Environ Biol 30:299–302.

Hartley-Whitaker J, Ainsworth G, Meharg AA (2001) Copper and arsenate-induced oxidative stress in Holcus lanatus L. clones with differential sensitivity. Plant Cell Environ 24:713–722.

Hashem A, Abd-Allah EF, Alqarawi AA, Huqail AAA, Egamberdieva D, Wirth S (2016) Alleviation of cadmium stress in Solanum lycopersicum L. by arbuscular mycorrhizal fungi via induction of acquired systemic tolerance. Saudi J Biol Sci 23:272–281. 10.1016/j.sjbs.2015.11.002

He Z, Shentu J, Yang X, Baligar VC, Zhang T, Stoffella PJ (2015) Heavy metal contamination of soils: sources, indicators, and assessment. J Environ Indicators 9:17–18.

Huang Y, He C, Shen C, Guo J, Mubeen S, Yuan J, Yang Z (2017) Toxicity of cadmium and its health risks from leafy vegetable consumption. J Funct Foods 8:1373–1401. 10.1039/c6fo01580h

Huang Y, Zu L, Zhang M, Yang T, Zhou M, Shi C, Shi F, Zhang W (2020) Tolerance and distribution of cadmium in an ornamental species Althaea rosea Cavan. Int J Phytoremediation 22:713–724. 10.1080/15226514.2019.1707771

Irfan M, Ahmad A, Hayat S (2014) Effect of cadmium on the growth and antioxidant enzymes in two varieties of Brassica juncea. Saudi J Bio Sci 21:125–131. 10.1016/j.sjbs.2013.08.001

Jamieson HE (2014) The legacy of arsenic contamination from mining and processing refractory gold ore at Giant Mine, Yellowknife, Northwest Territories, Canada. Rev Mineral Geochem 79:533–551. 10.2138/rmg.2014.79.12

Järup L, Akesson A (2009) Current status of cadmium as an environmental health problem. Toxicol Appl Pharmacol 238:201–208. 10.1016/j.taap.2009.04.020

Jomova K, Jenisova Z, Feszterova M, Baros S, Liska J, Hudecova D, Rhodes CJ, Valko M (2011) Arsenic: toxicity, oxidative stress and human disease. J Appl Toxicol 31:95–107. 10.1002/jat.1649

Juliana OO, Raymond WA, Tor-Anyiin TA, Dooshima TR (2019) Phytoremedation potential of Senna occidentalis to remove heavy metals from waste soil in Makurdi, Nigeria. Chem Mater 11:30–36. doi:10.7176/CMR/11-4-05

Kamunda C, Mathuthu M, Madhuku M (2016) Health risk assessment of heavy metals in soils from Witwatersrand Gold Mining Basin, South Africa. Int J Environ Res Public Health 13:663. 10.3390/ijerph13070663

Kulshrestha S, Dabral SK (2016) Bio-accumulation potential of wild plant Cassia Occidentalis L. grown on contaminated soil for Cd, Cr, Cu, Ni and Pb. JETIR 3:744–749.

Liu JG, Cai GL, Qian M, Wang D, Xu J, Yang J, Zhu Q (2007) Effect of Cd on the growth, dry matter accumulation and grain yield of different rice cultivars. J Sci Food Agric 87:1088–1095. 10.1002/jsfa.2816

Majumder B, Das S, Mukhopadhyay S, Biswas AK (2019) Identification of arsenic-tolerant and arsenic-sensitive rice (Oryza sativa L.) cultivars on the basis of arsenic accumulation assisted stress perception, morpho-biochemical responses, and alteration in genomic template stability. Protoplasma 256:193–211. doi: 10.1007/s00709-018-1290-5.

Marchiol L, Assolari S, Sacco P, Zerbi G (2004) Phytoextraction of heavy metals by canola (Brassica napus) and radish (Raphanus sativus) grown on multicontaminated soil. Environ Poll 132:21–27. 10.1016/j.envpol.2004.04.001

Marmiroli M, Imperiale D, Maestri E, Marmiroli N (2013) The response of Populus spp. to cadmium stress: Chemical, morphological and proteomics study. Chemosphere 93:1333–1344. 10.1016/j.chemosphere.2013.07.065

Martin SR, Llugany M, Barceló J, Poschenrieder C (2012) Cadmium exclusion a key factor in differential Cd-resistance in Thlaspi arvense ecotypes. Biol Plant 56:729–734. 10.1007/s10535-012-0056-8

McGrath SP, Zhao FJ (2003) Phytoextraction of metals and metalloids from contaminated soils. Curr Opin Biotech 14:227–282. 10.1016/S0958-1669(03)00060-0

Meharg AA, Macnaib MR (1990) An altered phosphate uptake system in arsenate tolerant Holcus lanatus L. New Phytol 116:29–35. 10.1111/j.1469-8137.1990.tb00507.x

Mellem JJ, Baijnath H, Odhav B (2012). Bioaccumulation of Cr, Hg, As, Pb, Cu, and Ni with the ability for hyperaccumulation by Amaranthus dubius. Afr J Agric Res 7:591–596. 10.5897/AJAR11.1486

Mishra S, Dubey RS (2006) Inhibition of ribonuclease and protease activities in arsenic exposed rice seedlings: Role of proline as enzyme protectant. J Plant Physiol 163:927–936. 10.1016/j.jplph.2005.08.003

Mrnka L, Kuchár M, Cieslarová Z, Matějka P, Száková J, Tlustoš P, Vosátka M (2012) Effects of endo- and ectomycorrhizal fungi on physiological parameters and heavy metals accumulation of two species from the family Salicaceae. Water Air Soil Pollut 223:399–410. 10.1007/s11270-011-0868-8

Muradoglu F, Gundogdu M, Ercisli S, Encu T, Balta F, Jaafar HZE, Zia-Ui-Haq M (2015) Cadmium toxicity affects chlorophyll a and b content, antioxidant enzyme activities and mineral nutrient accumulation in strawberry. Biol Res 48:11. 10.1186%2Fs40659-015-0001-3

Mukhtar S, Bhatti HN, Khalid M, Haq MAU, Shahzad SM (2010) Potential of sunflower (Helianthus annuus L.) for phytoremediation of nickel (Ni) and lead (Pb) contaminated water. Pak J Bot 42:4017–4026.

Niazi NK, Bibi I, Fatimah A, Shahid M, Javed MT, Wang H, Ok YS, Bashir S, Murtaza B, Saqib ZA (2017) Phosphate-assisted phytoremediation of arsenic by Brassica napus and Brassica juncea: Morphological and physiological response. Int J Phytoremed 19:670–678. 10.1080/15226514.2016.1278427.

Opdenakker K, Remans T, Keunen E, Vangronsveld J, Cuypers A (2012) Exposure of Arabidopsis thaliana to Cd or Cu excess leads to oxidative stress mediated alterations in MAPKinase transcript levels. Environ Exp Bot 83:53–61. 10.1016/j.envexpbot.2012.04.003

Ozbek K, Cebel N, Unver I (2014) Extractability and phytoavailability of cadmium in Cd-rich pedogenic soils. Turk J Agric For 38:70–79.

Petrovic D, Krivokapic S (2020) The Effect of Cu, Zn, Cd, and Pb Accumulation on Biochemical Parameters (Proline, Chlorophyll) in the Water Caltrop (Trapa natans L.), Lake Skadar, Montenegro. Plants (Basel) 9:1287. doi: 10.3390/plants9101287

Pietrini F, Iannelli MA, Pasqualini S, Massacci A (2003) Interaction of cadmium with glutathione and photosynthesis in developing leaves and chloroplasts of Phragmites australis (Cav.) Trin. ex strudel. Plant Physiol 133:829–837. 10.1104/pp.103.026518

Reeves PG, Chaney RL (2008) Bioavailability as an issue in risk assessment and management of food cadmium: A review. Sci Total Environ 398:13–19. 10.1016/j.scitotenv.2008.03.009

Sánchez-Viveros G, Ferrera-Cerrato R, Alarcón A (2011) Short-term effects of arsenate-induced toxicity on growth, chlorophyll and carotenoid contents, and total content of phenolic compounds of Azolla filiculoides. Water Air Soil Pollut 217:455–462. 10.1007/s11270-010-0600-0

Shahid M, Dumat C, Khalid S, Niazi NK, Antunes PMC (2017) Cadmium bioavailability, uptake, toxicity and detoxification in soil-plant system. Rev Environ Contam Toxicol 241:73–137. 10.1007/398_2016_8

Shanker AK, Cervantes C, Loza-Tavera H, Avudainayagam S (2005) Chromium toxicity in plants. Environ Int 31:739–753. 10.1016/j.envint.2005.02.003

Shi X, Wang S, Sun H, Chen Y, Wang D, Pan H, Zou Y, Liu J, Zheng L, Zhao X, Jiang Z (2017) Comparative of Quercus spp. and Salix spp. for phytoremediation of Pb/Zn mine tailings. Environ Sci Pollut Res Int 24:3400–3411. doi:10.1007/s11356-016-7979-0.

Silva-Ortega CO, Ochoa-Alfaro AE, Reyes-Agüerob JA, Aguado-Santacruz GA, Jimenez-Bremont JF (2008) Salt stress increases the expression of P5CS gene and induces proline accumulation in cactus pear. Plant Physiol Biochem 46:82–92. 10.1016/j.plaphy.2007.10.011

Silva JRR, Fernandes AR, Silva Junior ML, Santos CRC, Lobato AKS (2018) Tolerance mechanisms in Cassia alata exposed to cadmium toxicity – potential use for phytoremediation. Photosynthetica 56:495. 10.1007/s11099-017-0698-z

Singh HP, Batish DR, Kohli RK, Arora K (2007) Arsenic-induced root growth inhibition in mung bean (Phaseolus aureus Roxb.) is due to oxidative stress resulting from enhanced lipid peroxidation. Plant Growth Regul 53:65–73. 10.1007/s10725-007-9205-z

Singh S, Sinha S (2005) Accumulation of metals and its effects in (Brassica juncea L) Czern (cv Rohini) grown on various amendments of tannery waste. Ecotox Environ Safe 62:118–127. 10.1016/j.ecoenv.2004.12.026

Siripornadulsil S, Traina S, Verma DP, Sayre RT (2002) Molecular mechanisms of proline-mediated tolerance to toxic heavy metals in transgenic microalgae. Plant Cell 14:2837–2847. 10.1105/tpc.004853

Tiwari S, Gajbhiye T, Pandey SK (2016) Effect of Cadmium on morphological and biochemical constituents of Cassia species. JECET 5:379–390.

Ullah S, Khan J, Hayat K, Abdelfattah Elateeq A, Salam U, Yu B, Ma Y, Wang H, Tang ZH (2020) Comparative study of growth, cadmium accumulation and tolerance of three chickpea (Cicer arietinum L.) cultivars. Plants 9:310. 10.3390/plants9030310

U.S. EPA (2001) Method 200.7: Determination of Metals and Trace Elements in Water and Wastes by Inductively Coupled Plasma-Atomic Emission Spectrometry. Revision 5.1. Cincinnati (OH): USEPA-ICP Users Group.

Velikova V, Yondanov I, Edreva A (2000) Oxidative stress and some antioxidant system in acid rain treated bean plants: protective role of exogenous polyamines. J Plant Sci 151:59–66.

Verbruggen N, Hermans C (2008) Proline accumulation in plants: a review. Amino Acids 35:753–759. 10.1007/s00726-008-0061-6

Wu JT, Hsieh MT, Kow LC (1998) Role of proline accumulation in response to toxic copper in Chlorella sp. (Chlorophyceae) cells. J Phycol 31:113–117. 10.1046/j.1529-8817.1998.340113.x

Wuana R, Okieimen FE (2011) Heavy metals in contaminated soils: a review of sources, chemistry, risks and best available strategies for remediation. ISRN Ecol 2011:1–20. 10.5402/2011/402647

Yadav JP, Arya V, Yadav S, Panghal M, Kumar S, Dhankhar S (2010) Cassia occidentalis L.: a review on its ethnobotany, phytochemical and pharmacological profile. Fitoterapia 81:223–30. doi: 10.1016/j.fitote.2009.09.008.

Yoon J, Cao X, Zhou Q, Ma LQ (2006) Accumulation of Pb, Cu, and Zn in native plants growing on a contaminated Florida site. Sci Total Environ 368:456–464. 10.1016/j.scitotenv.2006.01.016

Zauro SA, Lawal AM, Umar KJ, Sani YM, Abubakar I (2015) Assessment of selected heavy metals in soil and Cassia Occidentalis in Rural area of Jega local government, Kebbi State, Nigeria. IJSRP 5:1–6.

Zhang H, Xu Z, Guo K, Huo Y, He G, Sun H, Guan Y, Xu N, Yang W, Sun G (2020) Toxic effects of heavy metal Cd and Zn on chlorophyll, carotenoid metabolism and photosynthetic function in tobacco leaves revealed by physiological and proteomics analysis. Ecotoxicol Environ Saf 202:110856. doi: 10.1016/j.ecoenv.2020.110856.

